# Effects of hydrological change in fire-prone wetland vegetation: an empirical simulation

**DOI:** 10.1101/2022.06.17.496658

**Authors:** Tanya J. Mason, Gordana C. Popovic, Maeve McGillycuddy, David A. Keith

**Affiliations:** Centre for Ecosystem Science, UNSW, Sydney, Australia; Department of Planning and Environment, New South Wales, Australia; Stats Central, Mark Wainwright Analytical Centre, UNSW Sydney, Australia

## Abstract

Upland swamps are peat-accumulating, groundwater-dependent and fire-prone wetland ecosystems. Drying caused by anthropogenic processes such as underground mining, ditching and climate change may disrupt surface and groundwater flows effecting a bottom-up control on wetland expression. Fire is an endogenous, recurring disturbance that drives a top-down consumptive force in many of these systems. When compounded with anthropogenic drying, fire may facilitate permanent community transitions. A dearth of ecological data and temporal lags have hampered our ability to predict risks associated with multiple disturbances in wetland plant communities. We collected intact wetland mesocosms from valley floors and lower slopes of four undisturbed swamp sites. We transferred the mesocosms to a glasshouse and established three different soil moisture availability levels to simulate wetland drainage. After 20 months of the hydrological treatment, we simulated a fire event by sequentially applying biomass removal (clipping), heat and smoke to half of the mesocosms. We monitored species biomass, richness and composition over a 3.5-year time frame. We found evidence of a temporal lag in biomass response to low water availability and synergistic hydrological and fire effects on species richness. In unburnt conditions, richness declined with low water availability but was maintained under high and medium water availability. After simulated fire in medium water availability, however, richness also declined and converged with depauperate low water mesocosm richness. Representation by many obligate swamp species declined in low compared with high water availability mesocosms over time, an effect that was amplified by the fire treatment.

**Synthesis:** Our evidence of lagged effects of hydrological change on wetland vegetation and compounding effects of fire should be considered in impact assessments, monitoring programs and ecosystem management to avoid irreversible wetland change in drying environments.

## Introduction

The relative contribution of bottom-up (resource availability) or top-down (consumer pressure) processes to community assembly has long focused ecological enquiry (e.g. Hunter and Price, 1992, Hairston et al., 1960, Bond, 2021). Bottom-up forces may include abiotic resource availability such as nutrients or water (Matson and Hunter, 1992) while top-down forces may include predator or herbivore activity (Kriegisch et al., 2016, Spiller and Schoener, 1990) or even fire activity (Bond and Keeley, 2005) as consumptive controls. Bottom-up and top-down forces may act simultaneously or with alternating patterns of dominance (Litzow and Ciannelli, 2007). Analysis of vegetation change when these forces interact may assist in understanding the mechanism of freshwater wetland community transitions under environmental change.

Hydrology is a sensitive and fundamental driver of freshwater wetland function and ecosystem service provision. It is therefore unsurprising that bottom-up controls have been considered dominant in wetland systems (Moore and Schmitz, 2021 and references therein). Inundation duration, depth and frequency influence root hypoxia, plant zonation and ultimately species composition (Blom and Voesenek, 1996, Campbell et al., 2016). Hydrological regulating and provisioning services including water purification, flood mitigation and water supply, along with carbon sequestration and biodiversity (Ramsar Convention on Wetlands, 2018) are important ecosystem services offered by wetlands. However, delivery of these services is threatened by drying and associated change to bottom-up resource availability. At the local or regional scale, underground mining and fracking extractive processes (Mason et al., 2021) and ditching (Lõhmus et al., 2015) or surface drainage (Woo and Young, 2005) affect water availability. At the global scale, climate change (particularly declining precipitation : evapotranspiration ratios), eutrophication and acid rain cause large-scale disruption of hydrological resources (Zedler and Kercher, 2005).

Alteration of water quantity through a change in flooding regimes may profoundly affect wetland expression by advantaging a new suite of species and driving community shifts along hydrological gradients (e.g. Zweig et al., 2020). However, if community reorganisation remains within thresholds for wetland function, redistribution of species along a hydrological gradient could represent an autonomous adaptation of an ecosystem while maintaining its function (Serrao-Neumann et al., 2016). Indeed, obligate wetland species have demonstrated considerable physiological tolerance to drying and may simply use flooded conditions as refugia from terrestrial competitors (Campbell et al., 2016, Blom and Voesenek, 1996).

Periodic imposition of top-down consumptive forces such as fire may “reset” wetland community status along divergent bidirectional succession trajectories towards either terrestrial or aquatic endpoints (Zweig and Kitchens, 2009). Fire consumes aboveground biomass and initiates recruitment through heat, smoke or light stimuli and enhanced resource availability (Whelan, 1995). The degree to which fire mediates a successional reset depends on hydrologic and vegetation legacies within the community (see Zweig and Kitchens, 2009 for a related discussion). If fire occurs when a wetland is within natural hydrological thresholds, the pre-fire community may re-establish (e.g. Clarkson, 1997, Flores et al., 2016). However, when fire occurs during novel hydrological conditions, it may initiate alternative successional pathways (e.g. Sulwiński et al., 2020).

Interactions between top-down and bottom-up processes may influence both the trajectory and the rate of successional change (Zweig et al., 2020 and references therein). Hydrological drivers alone may initiate gradual community change: Lõhmus et al. (2015) found that stand replacement occurred over multiple decades following forestry drainage (Lõhmus et al., 2015). Merritt and Cooper (2000) found that channel vegetation responded to river flow regulation over similar time frames. However, an interaction with fire may have synergistic effects that accelerate such transitions. For example, Turetsky et al. (2011) found elevated combustion losses of soil carbon when wildfire and water table drainage interacted in boreal peatland, while de Oliveira et al. (2014) found synergistic effects of fire on flood-prone riparian species composition and stem density. These interactions between top-down and bottom-up processes can strongly affect vegetation and biodiversity outcomes (Hunter and Price, 1992, Naeem, 2008, Wilkinson and Sherratt, 2016). However, disentangling singular and synergistic effects and testing conceptual models require an empirical approach with realistic yet controlled conditions (Srivastava et al., 2004).

Upland swamps are climatically marginal ecosystems on uplands and plateaus of southeastern mainland Australia and Tasmania (Keith et al., 2014). These groundwater-dependent and peat-accumulating wetlands support mosaics of wet heath communities comprising sclerophyll shrubs and graminoids that range from Ti-tree thicket and Cyperoid heath in the wettest conditions to Restioid heath, Sedgeland and Banksia thicket in drier conditions (Keith and Myerscough, 1993). Within swamps, fire is a recurrent disturbance (Mooney et al., 2021) that shapes subcommunity distributions (Keith and Myerscough, 1993). Swamp-woodland mosaics are likely sensitive to climatic moisture and fire history (Keith et al., 2010) and recent longwall underground coal extraction has caused subsidence, diminishing water resources and homogenising of the hydrological gradient, effects that persist long after coal extraction (Mason et al., 2021). The recent 2019-20 major bushfire season in eastern Australia provided observational evidence that mined upland swamps experienced abiotic and biotic change relative to unmined swamps in the post-fire environment and exhibited symptoms of ecosystem collapse (Keith et al., 2022). While the results were consistent with top-down fire accelerating more gradual ecosystem responses to bottom-up hydrological change, controlled empirical manipulations are required. In the current study we postulated that: 1) hydrological change initiates a transition in which productivity and composition diverge from that in the groundwater-dependent community; and 2) that a subsequent fire event causes more rapid transformation to a novel state by resetting the community and enabling less hydrophilous species to gain dominance (Figure 1).

**Figure 1:**
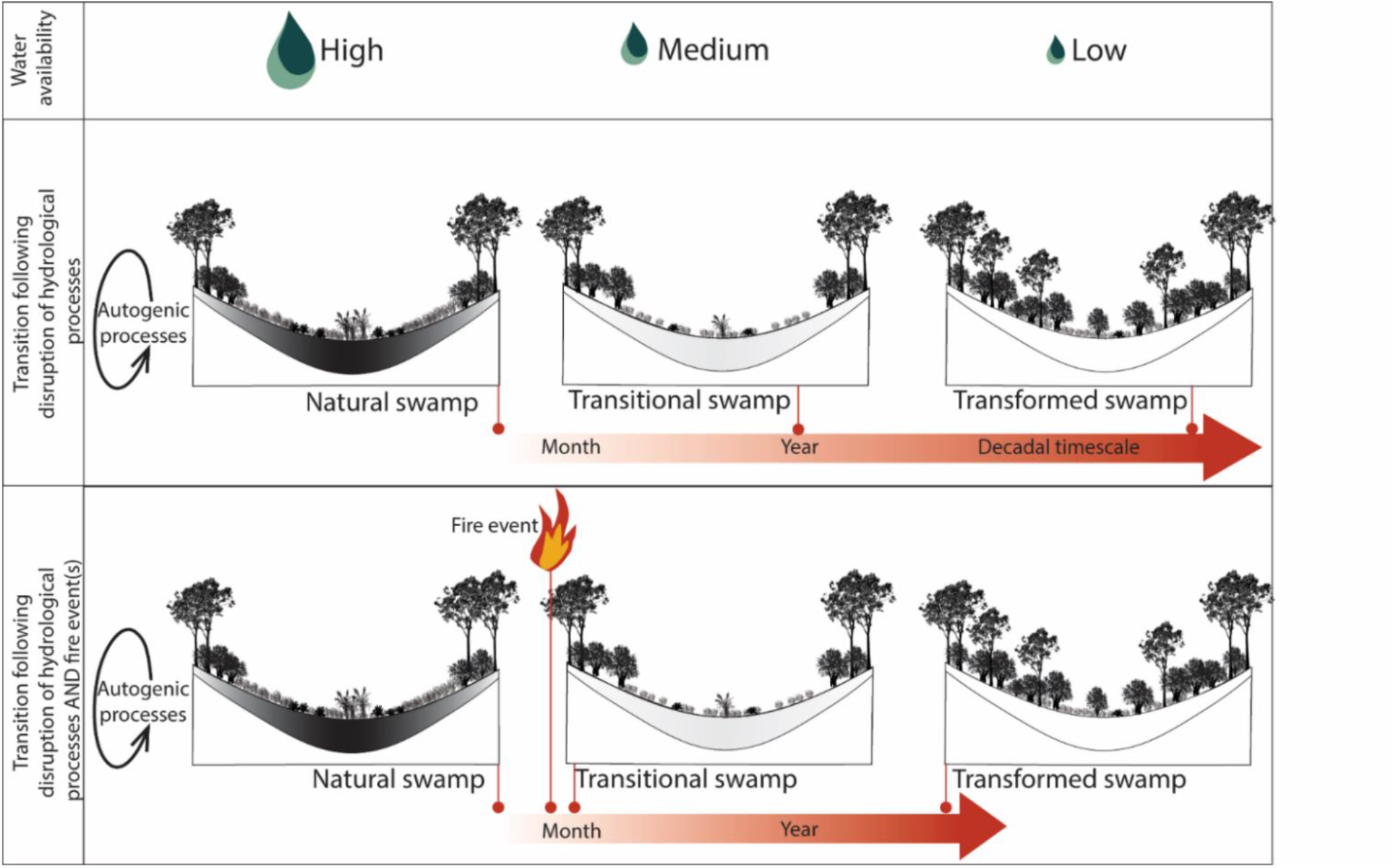
Conceptual model of swamp transition after changes to bottom-up (underground longwall mining) and top-down (periodic fire) processes over decadal time periods. Note that wet to dry soil conditions are illustrated with black (wet) gradation to white (dry) for the idealised hydrological gradient in the Natural swamp. Vegetation properties may differ among transformed swamps experiencing hydrological disruption alone vs. hydrological disruption and fire disturbance.

Here we experimentally assessed the influence of bottom-up and top-down forcing on upland swamp communities. We manipulated hydrological resource availability and fire occurrence in *ex situ* swamp mesocosms. Specifically, we asked: (1) Does reduced water availability affect wetland function and composition? (2) Is there evidence of a lagged response to bottom-up water resource availability? (3) Does fire interact with hydrological change by accelerating the lagged response in community function and composition? Comparing the importance of hydrological change and fire in determining biomass allocation and species composition will help to understand whether fire facilitates a transition of groundwater-dependent wetland communities to novel terrestrial ecosystems.

### Materials and Methods

#### Field collection

We collected intact mesocosms comprising above ground and root biomass along with soil material from four undisturbed upland swamp sites in the Sydney Basin during February and March 2017. Two sites were on the Woronora Plateau south of Sydney and two sites were in the Blue Mountains west of Sydney (Figure 2). Regional distinctions in species compositions have been recognised (Department of the Environment, 2014, Tozer et al., 2010).

**Figure 2:**
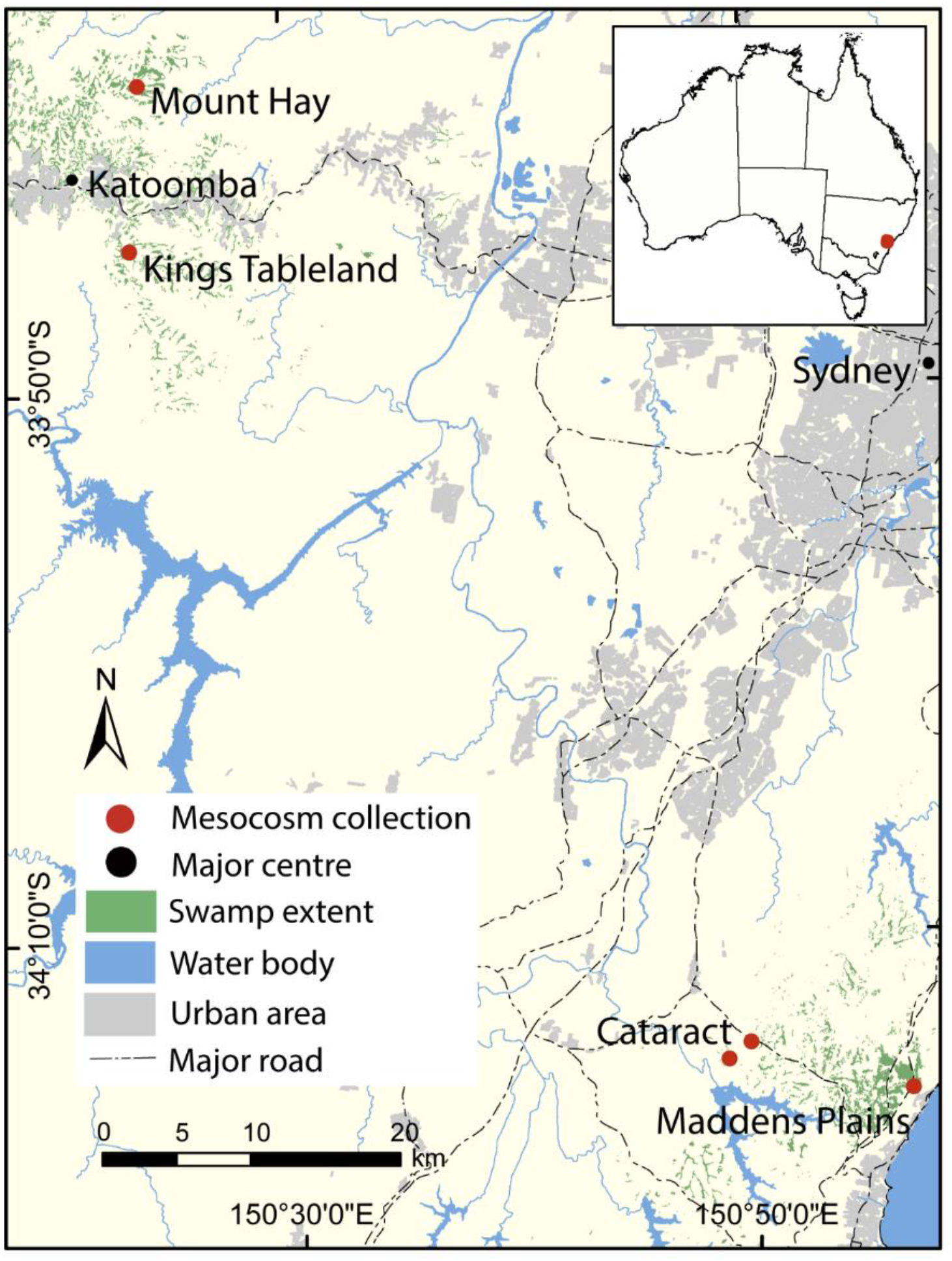
Location of mesocosm collection sites on the Woronora Plateau and Blue Mountains area. Inset top right shows location of study area (red) in Australia. Temperate Highland Peat Swamps on Sandstone layer obtained from Fryirs and Hose (2016). Note that transects at Cataract were at proximate swamps while transects at all other locations were at the same swamp.

#### Experimental procedure

We used a stratified, random approach to collect mesocosms along transects within both drainage zone (Ti-tree thicket) and mid slope (Cyperoid heath) plant communities. Mesocosms were positioned at least 0.5 m distant from each other and all swamp sites had been recently burnt (<20 months ago). In total, 248 mesocosms were collected (31 replicates x 2 plant communities x 4 sites) and their coordinates recorded. Mesocosms had a diameter of 150 mm and a depth of 250 mm and were housed in PVC sleeves which were hammered to ground level and extracted with trenching shovels. In order to minimise transplant shock, the above ground vegetation was clipped to approximately 10% of the original biomass and mesocosms were bagged, then transported to the glasshouse within 2 days of collection (Whalley and Brown, 1973). Mesocosms from each site were allowed to acclimatise for a month (28 days), with regular watering prior to treatment allocation.

The experimental setup was therefore staggered according to dates of field collection, commencing on the 24^th^ of March and completed on the 28^th^ of April 2017. We found that some mesocosms subsided in the PVC casing after collection. This was problematic as the treatment effect relied on water depth in the tubs. We packed the lower section of the subsided mesocosms with a 50:50 mix of river sand and peat moss to facilitate capillary action up the casing. All mesocosms were capped at the bottom with a permeable polyester fabric and affixed with cable ties. Our approach balanced tractability (replication and precise treatment application) with realism (initial species composition and nutrient regimes) (refer to Srivastava et al., 2004 for a related discussion, Naeem, 2001).

Mesocosms were randomly allocated to three water treatment levels and dispersed in tubs throughout the glasshouse. They were randomly relocated periodically within their treatment tubs across the glasshouse to minimise location and neighbour competition bias. We manipulated tub water levels to simulate different groundwater availability within mesocosms. We maintained water levels of treatment tubs (without top watering the mesocosms) to 70 mm, 155 mm and 240 mm below the mesocosm surface for the high, medium and low water level respectively. Initially we added water at weekly intervals (April 2017-December 2018). We increased the frequency of water addition (to biweekly) during summer (December to January) of 2018-19 to ensure emergence of seedlings immediately after fire simulation so their responses to water treatment could be tracked. We then added water at fortnightly (January 2019-November 2020) intervals to simulate ongoing drying of swamps above longwall mine paths (Mason et al., 2021). The same watering frequencies were applied across all water treatment levels. Volumetric soil water content (%) was monitored at interval using a Moisture Probe Meter (ICT International Pty Ltd MPM-160-B) with a 60 mm probe (average of three readings per mesocosm). Soil moisture values were initially within the range for the vadose zone of an undisturbed Cyperoid heath swamp (Mason et al., 2021). By the latter part of the experiment, the low water treatment yielded soil moisture values that overlapped the lower range for mined upland swamps (Keith et al., 2022).

After approximately 587 days (∼20 months) of the water treatment, we simulated a single fire event by sequentially clipping, heating and applying smoked water to half (124) of the mesocosms. We randomly selected mesocosms for burning treatment from each combination of water level and site. During October and November 2018, we clipped all biomass at ground level and bagged each species separately. We applied heat to the clipped soil surface using a propane burner (refer to Vesk et al., 2004 for a similar method). The burner was held approximately 10cm from the soil surface and the flame was constantly played across the mesocosm surface for three minutes. A pilot study indicated that 3 minutes was sufficient duration to provide temperatures > 60°C at 1 cm depth, and this was sufficient to break dormancy of some buried seed (Auld and Bradstock, 1996). We monitored heat penetration during the fire simulation by deploying a temperature datalogger (Thermochron ibutton, Maxim Integrated Products, Inc. San Jose, USA) at approximately 1 cm and 3 cm depth in each mesocosm. We used the maximum temperature (°C) recorded at each depth.

We generated smoked water following the method of Dixon et al. (1995) and using locally sourced leaf and twig material. The smoked water was applied to the clipped and heated mesocosms at a dilution of 1 part concentrate to 10 parts water (Read and Bellairs, 1999). Smoked water was applied to the soil surface 11 days after the heating component of the treatment. We used this approach to simulate a rain event after fire which would allow smoke particles to percolate through the soil and contact the seed bank. Unburnt mesocosms were top watered with tap water as a procedural control.

We monitored species composition at intervals throughout the experiment. We destructively harvested above ground biomass at two time points. Firstly, during the fire simulation and only for the fire-simulation mesocosms (∼ Day 587), and secondly at the conclusion of the experiment (∼ Day 1261) where above ground biomass was harvested for all mesocosms. In both cases, individuals were clipped at ground level, bagged, oven dried at 60°C for at least three days and weighed (AND GR-202 Series Balance ± 1.0 mg).

#### Data analysis

To examine the responses of above ground live biomass (measured at two time points) and species richness (measured at nine time points), we fitted linear mixed models with main effects and two-way interactions of time (categorical with 2 levels), water treatment (3 levels) and fire treatment (2 levels), controlling for fixed effects of swamp and vegetation type, and a random effect for mesocosm (to model dependence over time). To examine the responses of species composition, we used presence data and applied a multivariate linear mixed model, with a binomial distribution. The model used the same time points as the species richness linear model. We used a reduced rank correlation structure to model dependence between species (McGillycuddy et al. in prep). The model had fixed effects for time (linear), water treatment and fire treatment and their two-way interactions, and main effects of swamp and vegetation type. It also had (by species) random slopes corresponding to all the fixed effects, to allow species to respond differently to the treatment and environmental variables, as well as a mesocosm random effect to model dependence over time. Time was represented as a linear variable in the multivariate model as presence/absence data did not have enough information to estimate effects for each of the nine time points. The multivariate model was an extension of a generalised linear latent variable model (Niku et al., 2019) to allow additional random effects. This model allowed us to distinguish between (a) overall (uniform across species) trends in presence/absence by looking at fixed effects and (b) compositional changes - where species with atypical presence probabilities contributed to the composition effect. A treatment effect confidence interval that did not bracket zero suggested evidence of a treatment effect for that species. In addition, if it did not bracket the fixed effect value, then this species differed from the average treatment effect for all species, and therefore contributed to a compositional effect.

In all models, we examined two effects of interest: (a) time x water treatment interaction, to compare trends over time between water treatments, and (b) the water treatment x fire treatment interaction, to examine synergistic effects of water and fire treatments. In addition, using the multivariate model above, we fitted fourth corner models (Brown et al., 2014) by adding an interaction with species traits to determine if compositional changes were explained by two species traits, hydrological niche and fire response. Hydrological niches were assigned to species as *non-hydrophile* (no specific water demand in the substrate), *moderate hydrophile* (intermittently moist substrate) or *strong hydrophile* (permanently moist substrate) requirements. Fire responses were assigned to species as resprouter, killed, variable or unknown. Hydrological niches and fire responses were assigned to species based on Benson and McDougall (1993 -2002, 2005).

To verify the water treatment, we fitted a linear mixed model to soil moisture measured at 15 time points with water treatment (3 levels) and fire treatment (2 levels), as well as fixed effects of swamp and vegetation type, and a random effect for mesocosm (to model dependence over time). To verify the fire treatment, we fitted a model to mean maximum soil temperatures measured during the fire treatment. We included depth within soil profile (1cm or 3cm), water treatment and their interaction as fixed effects, and a random effect of mesocosm.

For all models, residuals were checked, and the response was transformed, or distribution changed, as necessary to meet model assumptions. The final model for biomass used log (biomass +1) as the response. For biomass and richness, we compared changes by treatment between two time points when data were mutually available: immediately prior to the fire treatment and at conclusion of the experiment. We adjusted for multiple pairwise tests using the multivariate t distribution. For multispecies models, we compared effects across species and traits, and did not control for multiple testing.

Most analyses used the glmmTMB package (Brooks et al., 2017) in R (R Core Team, 2021), with further analysis using the lme4 (Bates et al., 2015) and emmeans (Lenth, 2022) packages. We used lme4 when it interacted better with emmeans for rank deficient models (due to an unbalanced design for the fire treatment).

## Results

### Hydrological and fire treatment effects on soil moisture and temperature

We found very strong evidence that soil moisture levels differed as expected among the three water treatment levels (Figure 3). Low water mesocosms dried more quickly than either medium or high water mesocosms (water treatment : time interaction *F*30, 3674 = 19.070; *P* < 0.001, Figure 3). High water mesocosms also experienced reduction in soil moisture across the experiment, presumably as shoot and root growth progressed after initial clipping. Burnt mesocosms had an average 6.89% (95% CI: 5.49 - 8.29) higher soil moisture per volume than unburnt mesocosms, suggesting greater transpiration in unburnt compared with burnt mesocosms.

**Figure 3:**
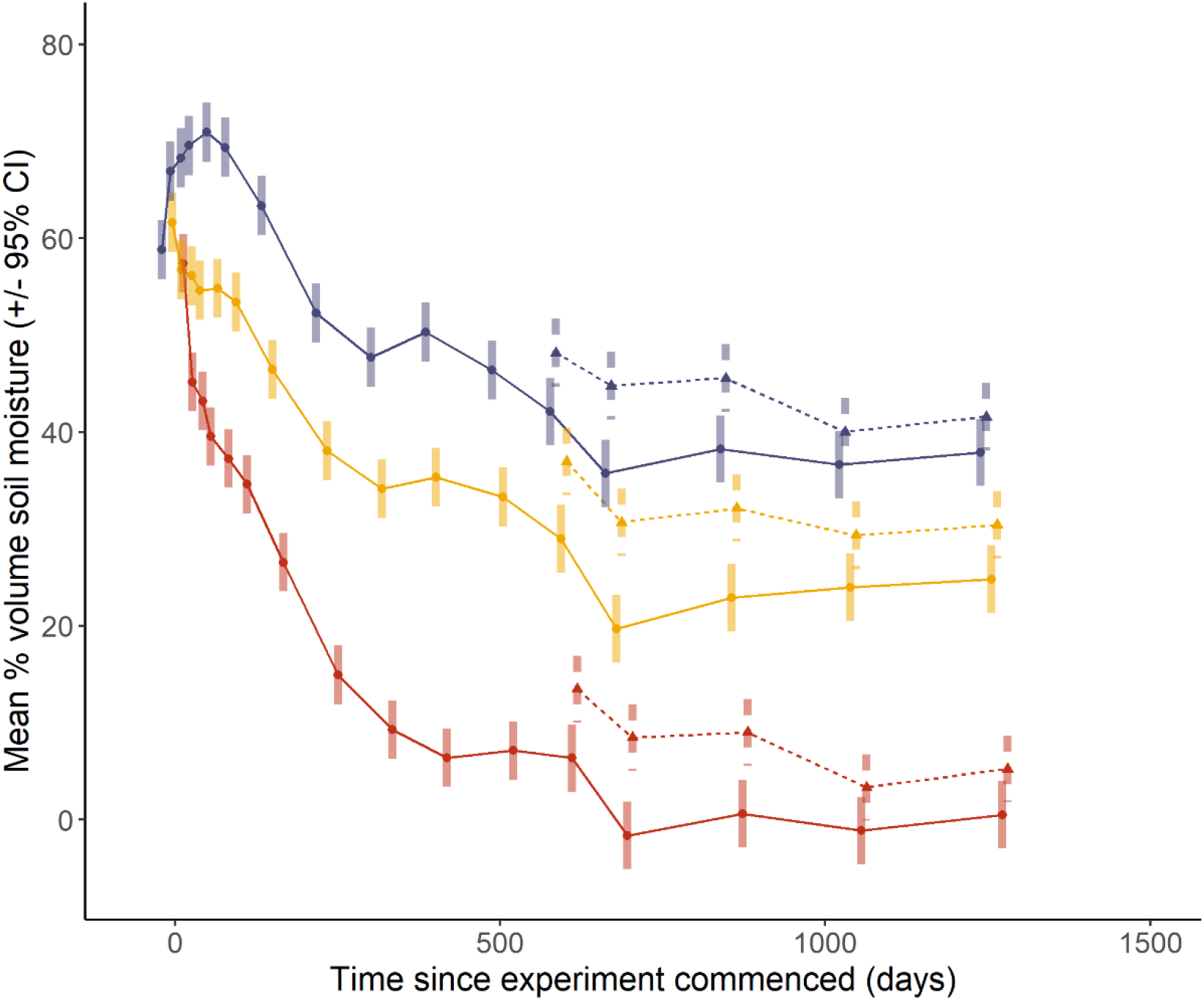
Estimated marginal means (± 95% confidence intervals) for mesocosm soil moisture (% volume) over time (days since commencement). Treatment levels are high 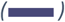 medium 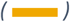 and low 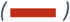 water availability and unburnt 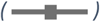 and burnt 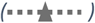 fire treatment levels.

Among burnt mesocosms, we found very strong evidence of differences in maximum soil temperature during the fire simulation. Temperatures depended on water availability and depth within the soil profile (water treatment : depth in soil profile interaction; χ^2^_2_ = 20.858; *P* < 0.001, Figure 4). Contrast analyses revealed that at each soil profile depth, low water mesocosms experienced hotter maximum temperatures when compared with medium and high water mesocosms. There was no evidence of differences in maximum temperatures for medium and high water mesocosms (Figure 4). The mean maximum temperature was 1.966 (95% CI: 1.617 -- 2.389) times higher in low water mesocosms than high water mesocosms at 1cm, and 1.461 (95% CI: 1.195 -- 1.786) times higher at 3cm depth. The mean maximum temperature was 1.670 (95% CI: 1.377 -- 2.024) times higher in low water mesocosms than medium water mesocosms at 1cm, and 1.333 (95% CI: 1.094 -- 1.625) times higher at 3cm depth.

**Figure 4:**
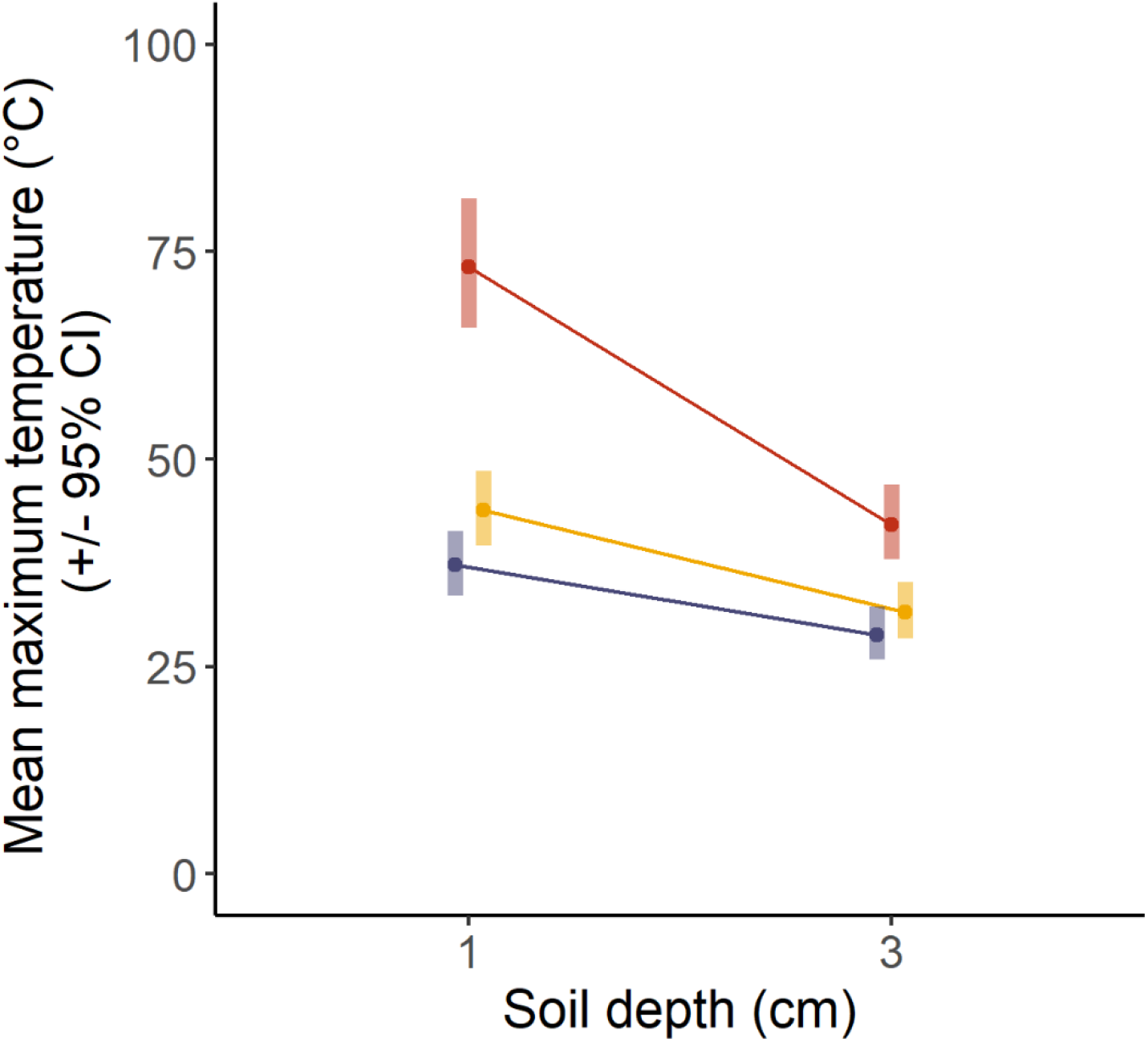
Estimated marginal means (± 95% confidence intervals) for maximum temperature (°C) at 1 cm and 3 cm soil profile depth in burnt mesocosms during fire simulation (Day 587 of experiment). Treatment levels are high 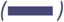 medium 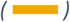 and low 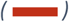 water availability.

### Hydrological and fire effects on above ground biomass

We found strong evidence that water treatment influenced changes in biomass over time (*F*2, 343 = 5.804; *P* = 0.003; Figure 5). Planned contrasts showed strong evidence (*t*341 = 3.093, *P* = 0.006) that differences in biomass between unburnt low and high water mesocosms increased over time: biomass differences between high and low water mesocosms more than doubled (2.183 (95% CI: 1.205 – 3.955)) between Day 587 and Day 1261 of the experiment. Similarly, differences in biomass between unburnt low and medium water mesocosms more than doubled between Day 587 and Day 1261 (*t*341 = 2.771, *P* = 0.016; estimate = 2.013; 95% CI: 1.111–3.646), but there was no evidence of differences in biomass changes between high and medium water mesocosms (*t*341 = 0.320, *P* = 0.945; estimate = 1.085; 95% CI: 0.597 – 1.971). So above-ground biomass showed a lagged response to low water availability. We did not find any evidence of an interaction between fire and water treatments (*F*2, 344 = 0.600; *P* = 0.549), and therefore no evidence of a synergistic effect of fire and water on above ground biomass (Figure 5).

**Figure 5:**
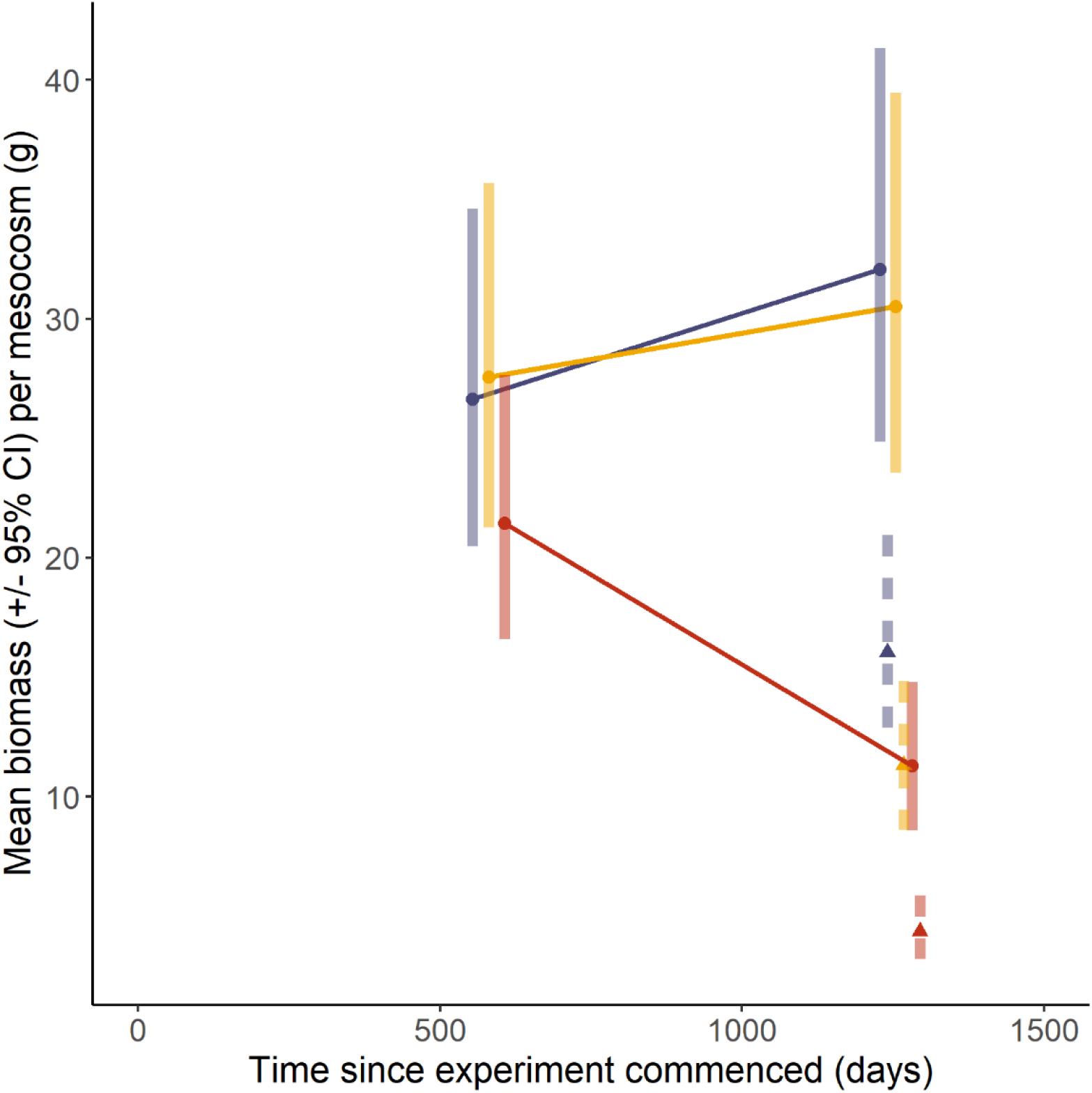
Estimated marginal means (± 95% confidence intervals) for above ground biomass (g) per mesocosm at Day 587 and 1261 since commencement of the experiment. Treatment levels are high 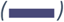 medium 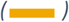 and low 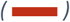 water availability and unburnt 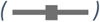 and burnt 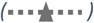 fire treatment levels. Note that biomass was collected immediately prior to the fire simulation and again at the end of the experiment.

### Hydrological and fire effects on species richness

We found very strong evidence that changes in richness over time differed by water treatment (*F*16,1960 = 9.878, *P* < 0.001) for unburnt mesocosms. Species richness in low water mesocosms diverged below richness in high water (H vs. L by time interaction t1974 = 6.027, *P* < 0.001; estimate = 1.635; 95% CI: 0.933 – 2.337) and medium water (M vs. L by time interaction t1974 = 4.943, *P* < 0.001; estimate = 1.344; 95% CI: 0.640 – 2.047) mesocosms. However, there was no evidence of divergence between high and medium water mesocosms (t1973 = 1.068; *P* = 0.759; estimate = 0.291; 95% CI: - 0.414 - 0.995, Figure 6). We found very strong evidence of an interaction between fire and water treatments - hence a synergistic effect of fire and water on richness (*F*2, 2044 = 18.801; *P* < 0.001; Figure 6). Contrasts showed very strong evidence of convergence between medium and low water availability after the fire treatment (burnt mesocosms) compared to the unburnt treatment level (t2044 = -4.829, *P* < 0.001; estimate = -1.160; 95% CI: -1.782 - -0.539), and divergence between high and medium water availability after fire (t2044 = 5.693, P < 0.001; estimate = 1.376; 95% CI: 0.751 – 2.001) compared to the unburnt treatment level. There was no evidence of change in the difference between richness for high and low water availability with fire (t2044 = 0.898, *P* < 0.854; estimate = 0.216; 95% CI: -0.406 – 0.838). In summary, the fire treatment accelerated species richness declines in medium water mesocosms, but not high water mesocosms. Richness declined in low water mesocosms independently of the fire treatment, but declines were not evident until nearly 2 years after the water treatment was implemented (Figure 6).

**Figure 6:**
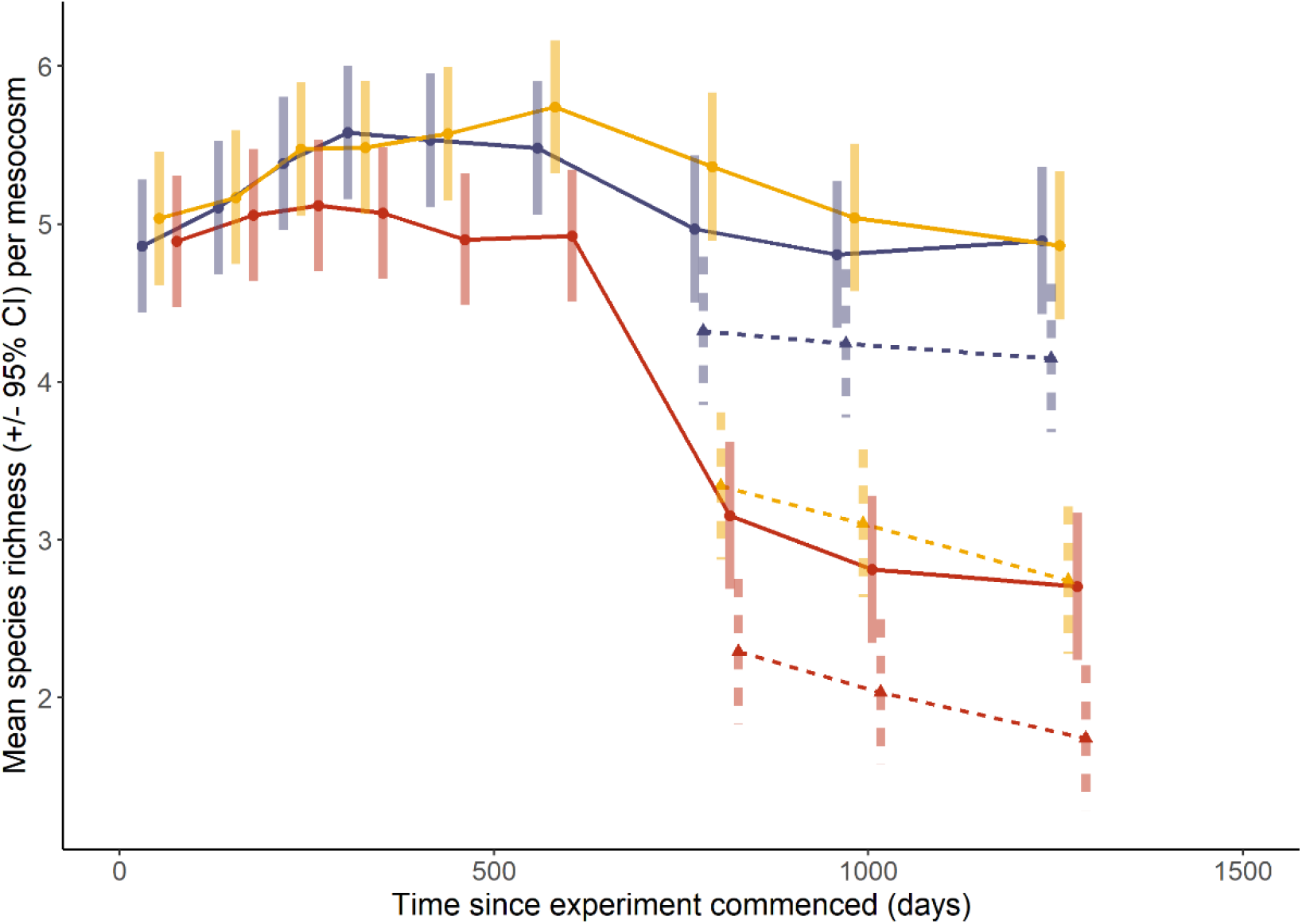
Estimated marginal means (± 95% confidence intervals) for species richness over time (days since commencement) in treatment mesocosms. Treatment levels are high 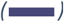 medium 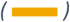 and low 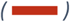 water availability and unburnt 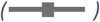 and burnt 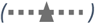 fire treatment levels.

### Hydrological effects on species composition and occurrence

Consistent with a reduction in richness, the odds of representation (i.e. presence) across species reduced 66% (95% CI: 49% - 78%) more in the low water than the high water mesocosms over the experimental period. This effect was driven by twenty-three species (Figure 7 (a) species in brown). The reduction of species representation in low water compared with high water availability mesocosms was not uniform across species, with two species experiencing particularly large reductions, and two species exhibiting smaller than average reductions (Figure 8 (a)). These variable responses drove the compositional change (χ^2^_2_ = 14.335; *P* < 0.001). We found evidence that some of this compositional change was explained by reduced representation by hydrophile species (χ^2^_12_ = 24.638; *P* = 0.017), however we did not identify particular pairwise differences between species with different hydrological niches: moderate and strong hydrophile species underwent similar reductions when exposed to low water availability, and non-hydrophile species had large standard errors (Appendix 1). There was no evidence of difference in average representation between medium and high water availability (OR = 0.89; 95% CI: 0.68 - 1.17) (Appendix 1).

**Figure 7:**
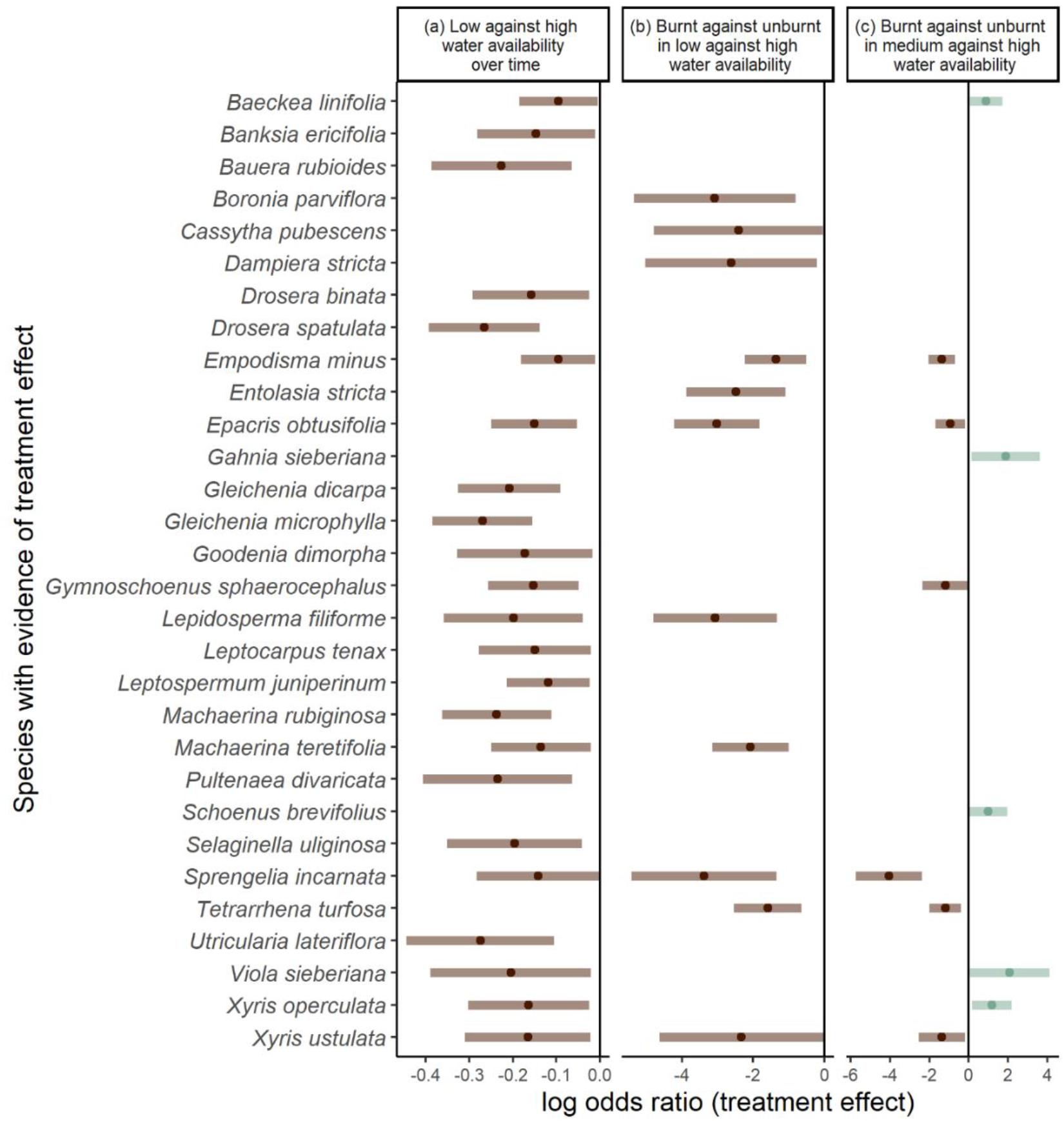
Treatment effect (log odds ratio ± 95% confidence interval) for species exhibiting (a) a change over time or (b-c) synergistic water and fire effect. Species shown in brown 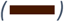 had large reductions in representation. Species shown in teal 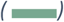 had smaller than average reductions or increases in representation.

**Figure 8:**
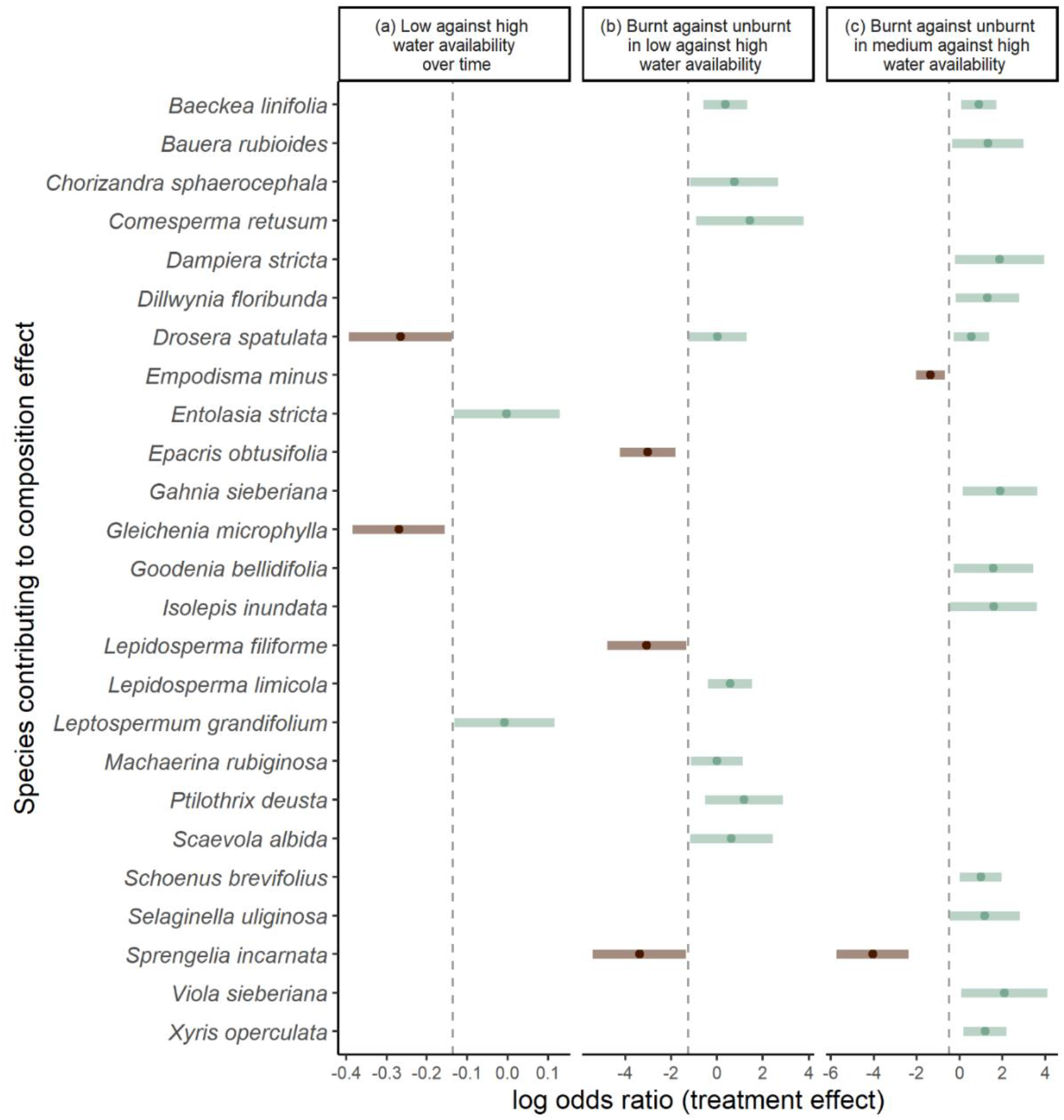
Treatment effect (log odds ratio ± 95% confidence interval) for species whose compositional contribution changed (a) over time or (b-c) with water and fire effects. Species shown in brown 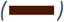 had large reductions in representation. Species shown in teal 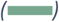 had smaller than average reductions or increases in representation.

Similarly to our species richness results, we found strong evidence of synergistic effects of water availability and fire disturbance on species composition. Burnt mesocosms had a larger difference in representation between high and low water availability than unburnt mesocosms (OR = 3.51, 95% CI: 1.67 – 7.39), while we did not find evidence of a similar difference for burnt *vs*. unburnt treatment levels in high and medium water availability mesocosms (OR = 1.62, 95% CI: 0.89 – 2.96). The interaction was driven by 17 species (Figure 7 (b-c) species in brown). The reduction was not uniform across all species (χ^2^_2_ = 78.773; *P* < 0.001): five species experienced particularly large reductions (Figure 8 (b-c) species in brown), and 20 had smaller than average reductions or increases (Figure 8 (c) species in teal) relative to unburnt mesocosms. We found no evidence (χ^2^_36_ = 33.914; *P* = 0.568) that this compositional change could be explained by fire traits (Appendix 2).

## Discussion

Wetland function was detrimentally affected by disruption of water resource availability. The rate and magnitude of that effect was influenced by fire-mediated top-down regulation. These experimental results support our conceptual model of swamp transition (Figure 1). We found very strong evidence that water availability is a bottom-up driver of swamp vegetation structure. Above ground biomass, species richness and composition were all detrimentally affected by low water availability. Persistent reduction in water availability will drive a loss of species and primary productivity within swamp communities. Over the experimental timeframe of 3.5 years, we observed a lagged response of above-ground biomass and species richness to hydrological change. By the end of the experiment however, reduced water availability had caused declines in primary productivity (biomass) and biodiversity value (species richness and representation).

Strongly hydrophilic species appeared to have lower representation in the low than high water mesocosms. But non-hydrophile species did not show evidence of differentiation in the low vs. high water availability mesocosms. In addition, species disproportionately adversely affected by low water availability were often obligate wetland species (e.g. *Glechenia microphylla*) while many species less adversely affected than average were facultative wetland species (e.g. *Entolasia stricta*). These emerging compositional patterns may signal increased terrestrialisation of swamp vegetation as mining-induced wetland drying continues.

We found that fire disturbance compounded hydrological change and accelerated swamp transition, corroborating our conceptual model and field-based observations (Keith et al., 2022). By the end of the experiment, differences in species compositions were greater under water stress for burnt than unburnt mesocosms. Post-fire weather conditions are among the most important stochastic environmental factors affecting vegetation dynamics in Australian ecosystems (Keith et al., 2002) such as Cyperoid heath and Ti-tree thicket. Recruitment may fail when low rainfall follows a fire event (Keith et al., 2002). In our case, low water availability appeared to limit regeneration of both resprouts and seedling recruits after the fire treatment. When combined, hydrological disruption and fire acted synergistically, with adverse effects on post-fire species richness and composition. While swamp communities may not immediately respond to changed water availability (evidenced by temporal lags in biomass and species richness in unburnt mesocosms), recurrent fire is likely to accentuate community change.

Our experiment simulated a low-intensity fire (Auld, 1986). Notably, low water mesocosms most effectively conducted heat down the soil profile. A study by Wright and Clarke (2008) on fire effects in central Australian spinifex communities also reported a strong effect of soil moisture on temperature profiles with high soil moisture strongly reducing soil heating during fire. Consumption of a large mass of ground fuels during fire is likely to generate hotter soil temperatures than those recorded during our fire simulation, and this may initiate smouldering substrate fires that cause elevated mortality of roots, seedbanks and rhizomes (Rein et al., 2008) in swamp sediments with low moisture content (Prior et al., 2020).

Fire stimulates plant emergence from seedbanks and underground organs (Keith, 2012). However, the interplay of fire and hydrology on biotic components of the soil profile may be complex. Tangney et al. (2018) identified a potentially countervailing phenomenon where elevated seed moisture content facilitated lower lethal temperature thresholds when compared with drier seed: when seeds and soil are moist, seed mortality increases. In our experiment, maximum soil temperatures for high and medium water mesocosms were mostly below temperature thresholds (50-180°C) used by Tangney et al. (2018). We therefore conclude that high and medium water levels in our experiment buffered against elevated soil temperatures and avoided risks of seed mortality (while probably also moderating the stimulative effect of fire on species richness).

After the fire simulation, we attributed higher soil moisture of burnt compared with unburnt mesocosms to lower above ground biomass and consequently lower transpiration rates in burnt mesocosms. Despite unconsumed water resources and potential stimulative fire effects, low water mesocosms when burnt, did not support higher species richness or greater representation of swamp species compared with low water availability unburnt mesocosms. Our experimental design could not identify whether soil temperatures increased seed mortality or whether low soil moisture inhibited seed germination and seedling survival in low water mesocosms. However, we have shown that multiple disturbance selectively filtered swamp species and affected their long-term representation. These environmental changes and losses of diversity may ultimately lead to ecosystem collapse, where ecosystem function and identity are transformed (Keith et al., 2022).

By extracting mesocosms and exposing them to controlled conditions in the glasshouse, we closely simulated field conditions and compositions. However, the realism of our experiment was constrained by restricting groundwater variability and excluding any simulation of precipitation. In addition, our simulation of intensified swamp drying, with a decrease in water addition from weekly to fortnightly frequency, occurred soon after the fire simulation. However, differences across water treatment levels were already evident prior to this change in the frequency of water addition. The character of vegetation transformed by the drying treatment in our mesocosm experiment likely differed from *in situ* vegetation of drying swamps: glasshouse conditions essentially precluded colonisation from the regional species pool, including terrestrial species of the surrounding woodland matrix. In the absence of colonisation opportunities, and over our truncated timeline, we are unable to identify outcomes of the community transition. Natural experiments in mined and unmined swamps (e.g. Keith et al., 2022) provide some insights into the character of the derived communities that replace collapsed upland swamp ecosystems. Thus far, research indicates that swamps may either transition to terrestrial communities or remain as depauperate swamps comprising facultative rather than obligate swamp species (Keith et al., 2022).

### Conservation and management implications

Upland swamps of the Sydney Basin are substantively threatened by subsidence disturbance and disruption of groundwater following underground mining. While our empirical study was initially aimed at simulating the effects of underground mining, global climate change (where increased evapotranspiration and reduced precipitation occur) or deforestation and landuse change may similarly affect water budgets. Wherever long-term hydrological disruption of wetland ecosystems occurs, policymakers should expect declines in ecosystem function and biodiversity. When combined with recurrrent, endogenous disturbance such as fire, the impacts of hydrological change may be amplified and transition to alternative ecosystem states accelerated. Ultimately, the effects of hydrological disturbance cannot be comprehensively assessed by planning organisations without also considering recurrent disturbances - such as fire - in a landscape context.

Long-term field monitoring has indicated that, in the absence of exogenous hydrological change, plant communities in the wettest parts of swamps (Ti-tree thicket and Cyperoid heath) are more resistant to compositional change than those in drier parts of swamps (Restioid heath, Sedgeland and Banksia thicket) (Mason et al., 2017). Our current findings have shown that even plant communities in the wettest habitats are susceptible to transformation if hydrological thresholds are exceeded. Mining disturbance may entrain irreversible change in swamp vegetation and associated biota.

Upland swamps near the footprint of underground longwall mining (within a buffer zone ∼600m) may experience drying due to increased permeability of sandstone aquifers in the surface cracking zone (Watershed HydroGeo, 2022). Within the buffer zone, swamps that source water from a regional water table rather than relying on rainfall alone, may also experience drying or partial drying (David et al., 2017). We found that even under partial dewatering, species richness was disproportionately diminished after fire. Consequently, swamps within and near the longwall mine footprint may be vulnerable to transition when fire follows hydrological disturbance and mine-related subsidence impacts should not be assumed to be confined to the direct underground mining footprint.

Upland swamp conservation requires an understanding of both bottom-up and top-down regulation processes. Substantial reductions in soil moisture initiates ecosystem-level change. Longwall mining generates almost immediate hydrological disruption (Mason et al., 2021) followed by longer-term, lagged biotic change as measured in our simulation experiment and in the field (Keith et al., 2022). The challenge for policymakers is to assess impacts of disturbance when plant community responses are lagged, cumulative or chaotic (Milchunas and Lauenroth, 1995). Climate change, while a less abrupt process, may have similar long term effects on swamp hydrology (Keith et al., 2014). Under both regional and global hydrological change, positive feedbacks may establish: dry conditions create more favourable fire weather and more frequent fire disturbance. Synergistic hydrological and fire disturbance accelerate vegetation and ecosystem change and imperil swamp conservation. Planners and policy makers must be cognizant that once hydrological disturbance occurs, compounding physical and biotic change are likely unavoidable.

## Author contributions

Tanya Mason, David Keith and Gordana Popovic conceived the ideas and designed methodology; Tanya Mason collected the data; Gordana Popovic, Maeve McGillycuddy and Tanya Mason analysed the data; Tanya Mason led the writing of the manuscript. All authors contributed critically to the drafts and gave final approval for publication.

## Acknowledgements

We acknowledge the Dharawal and Dharug people, who are the traditional custodians of the land on which this study was conducted. We acknowledge field, glasshouse and laboratory assistance from Ava Chamberlain, Len Martin, Katy Wilkins, Chantel Benbow, Douglas Simpson, Ada Sanchez-Mercado and Christopher Simpson. We thank the NSW Department of Planning and Environment for access to their Lidcombe glasshouse facility. This project has been assisted by the NSW Government through its Environmental Trust (2018/SSC/0049) and Saving Our Species program (Conservation of threatened groundwater dependent ecosystems: understanding resilience and adaptive capacity to hydrological change). Fieldwork was conducted under DPE Scientific Licence SL101124. Data and R code used in the study are available at https://github.com/mmcgillycuddy/wetland_paper.

## Appendices

**Appendix 1:**
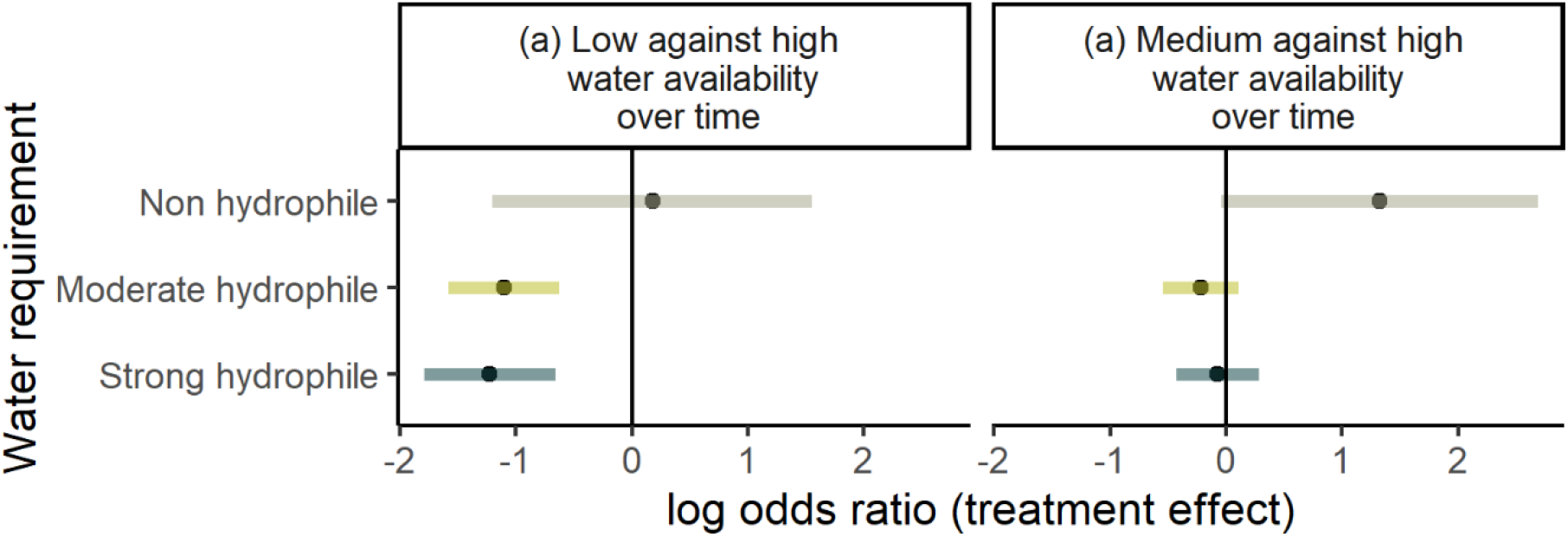
Water availability treatment effect (log odds ratio ± 95% confidence interval) on water requirement traits over time.

**Appendix 2:**
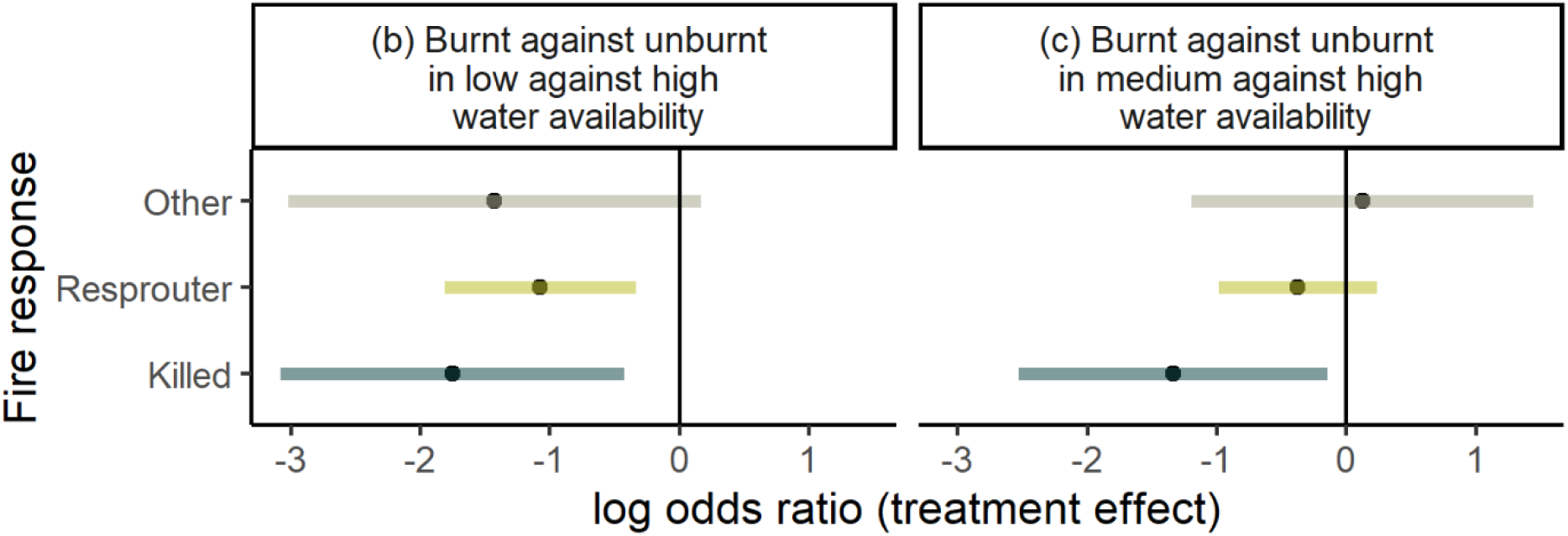
Fire treatment effect (log odds ratio ± 95% confidence interval) on fire response traits. “Other” species have either variable or unknown responses.

